# Probing mycelium mechanics and taste: The moist and fibrous signature of fungi steak

**DOI:** 10.1101/2025.04.17.649423

**Authors:** Thibault Vervenne, Skyler R. St. Pierre, Nele Famaey, Ellen Kuhl

## Abstract

Fungi-based meat is emerging as a promising class of nutritious, sustainable, and minimally processed biomaterials with the potential to complement or replace traditional animal and plant-based meats. However, its mechanical and sensory properties remain largely unknown. Here we characterize the quasi-static and dynamic mechanical behavior of fungi-based steak using multi-axial mechanical testing, rheology, and texture profile analysis. We find that the rate-independent response under quasi-static compression and shear is isotropic, while the rate-dependent response under dynamic compression is markedly anisotropic with stiffnesses and peak forces four times larger cross-plane than in-plane. Automated model discovery reveals that the exponential Demiray model best explains the rate-independent elastic response upon chewing in both directions. The rate-dependent directional stiffening can be linked, at least in part, to the high water content of mushroom root mycelium and to the restricted fluid flow at higher loading rates. Complementary sensory surveys reveal a strong correlation with mechanical metrics and suggest that we perceive the fungi-based steak as more moist, more viscous, and more fibrous than traditional animal- and plant-based meats. Taken together, our findings position fungi-based steak as an attractive, structurally equivalent, and sensorially compelling alternative protein source that is healthy for people and for the planet.

## 1. Introduction

The growing global demand for protein creates an urgent need for nutritious and sustainable alternatives to conventional animal meat [1]. Fungi-based proteins from the filamentous mushroom root offer a promising solution [2]. Mycelium is the vegetative lower part of fungi and the largest living organism on earth, that can occupy up to 10 km^2^ [3]. Nutritionally, mycelium is low in fat and rich in protein, fibers, and vitamins [4]; it contains all essential amino acids [5]; and it provides health benefits that enhance human immunity, decrease cholesterol levels, and improve the gut microbiome [6]. Structurally, mycelium forms a dense entangled network of branching micro-filaments called hyphae–an assembly of elongated cells, separated by cross walls called septa, all enclosed within the cell wall–resembling the fibrous microstructure of animal meat [7]. Environmentally, mycelium grows rapidly and robustly on low-cost agricultural substrates with minimal land, water, and energy use [8], and generates significantly lower greenhouse gas emissions than traditional animal agriculture [9]. These natural benefits position fungi-based meats as innovative next-generation proteins with the potential to address challenges in both human and planetary health.

Despite this promise, the use of mycelium as a functional biomaterial for food remains poorly understood. The fibrous, filamentous microstructure of mycelium is distinct from both animal and plant tissues [7], and its bulk mechanical properties have not been systematically characterized. While several studies have explored mycelium-based composites for packaging, insulation, and building materials [10], the rheological and textural behavior of edible mycelium-based products remains virtually unstudied. No standardized constitutive model exists for fungi-based meat, and little is known about how its mechanical response varies with the mode of deformation, and with the loading magnitude, rate, and direction. Without this understanding, it is challenging to rationally design fungi-based products that match the mechanical and sensory profiles of traditional animal meat [11].

In this study, we present a comprehensive biomechanical and sensory characterization of a commercially available fungi-based steak. We perform controlled compression and shear tests in both in-plane and cross-plane directions [12], quantify texture using double-compression texture profile analysis and oscillatory rheology [13], and discover the best fungi steak model and parameters using automated model discovery [14]. To explore to which extent these mechanical features are predictive of our sensory perception, we conduct a supplementary sensory survey. We invite sixteen participants to taste and score twelve sensory features, both in-plane and cross-plane, and systematically correlate these features to our mechanical tests [15]. Finally, we compare the performance of the fungi-based steak against eight common commercial meat products [11]. Our overarching question is: *Can fungi-based steak reliably match or exceed the key mechanical and sensory features of traditional animal- and plant-based meats?*

## 2. Methods

Throughout this study, we test the *fungi-based meat* Mushroom Root Classic Steak (Meati Foods, Boulder, CO) in two orthogonal directions, the *in-plane direction* of the steak parallel to the fiber direction, and the *cross-plane direction* of the steak perpendicular to the fiber direction. According to the ingredient list, the product contains, in descending order by weight, mushroom root (mycelium), and less than 2% of oat fiber, salt, fruit juice for color, vegetable juice for color, natural flavor, and lycopene for color, with mycelium making up 95% of the product. According to the nutrition label, one serving of one steak weighs 120 g and contains 1.5 g or 2% of the percentage daily value of total fat, 9 g or 3% of daily total carbohydrates, with 8 g or 29% of daily dietary fiber, 15 g or 30% of daily protein, and 200 mg or 9% of daily sodium.

We compare the fungi-based meat against three *animal-based meats*, Turkey Polska Kielbasa Sausage (Hillshire Farm, New London, WI), Spam Oven-Roasted Turkey (Spam, Hormel Foods Co, Austin, MN), and Classic Uncured Wieners (Oscar Mayer, Kraft Heinz Co, Chicago, IL), and five *plant-based meats*, Ham-Style Roast Tofurky (Tofurky, Hood River, OR) Vegan Frankfurter Sausage (Field Roast, Seattle, WA), Plant-Based Signature Stadium Hot Dog (Field Roast, Seattle, WA), Organic Tofu Extra Firm (House Foods, Garden Grove, CA), and Organic Firm Tofu (365 by Whole Foods, Austin, TX). We select these eight comparison products from our previous texture profile analysis and rheological tests [13], and our mechanical tests and sensory surveys [11].

### 2.1. Sample preparation

We use a biopsy punch to extract a total of *n* = 164 cylindrical samples of 8 mm diameter and 10 mm height, *n* = 82 *in-plane*, parallel to the fiber direction, and *n* = 82 *cross-plane*, perpendicular to the fiber direction. Figure 1 shows the sample preparation process.

**Figure 1:**
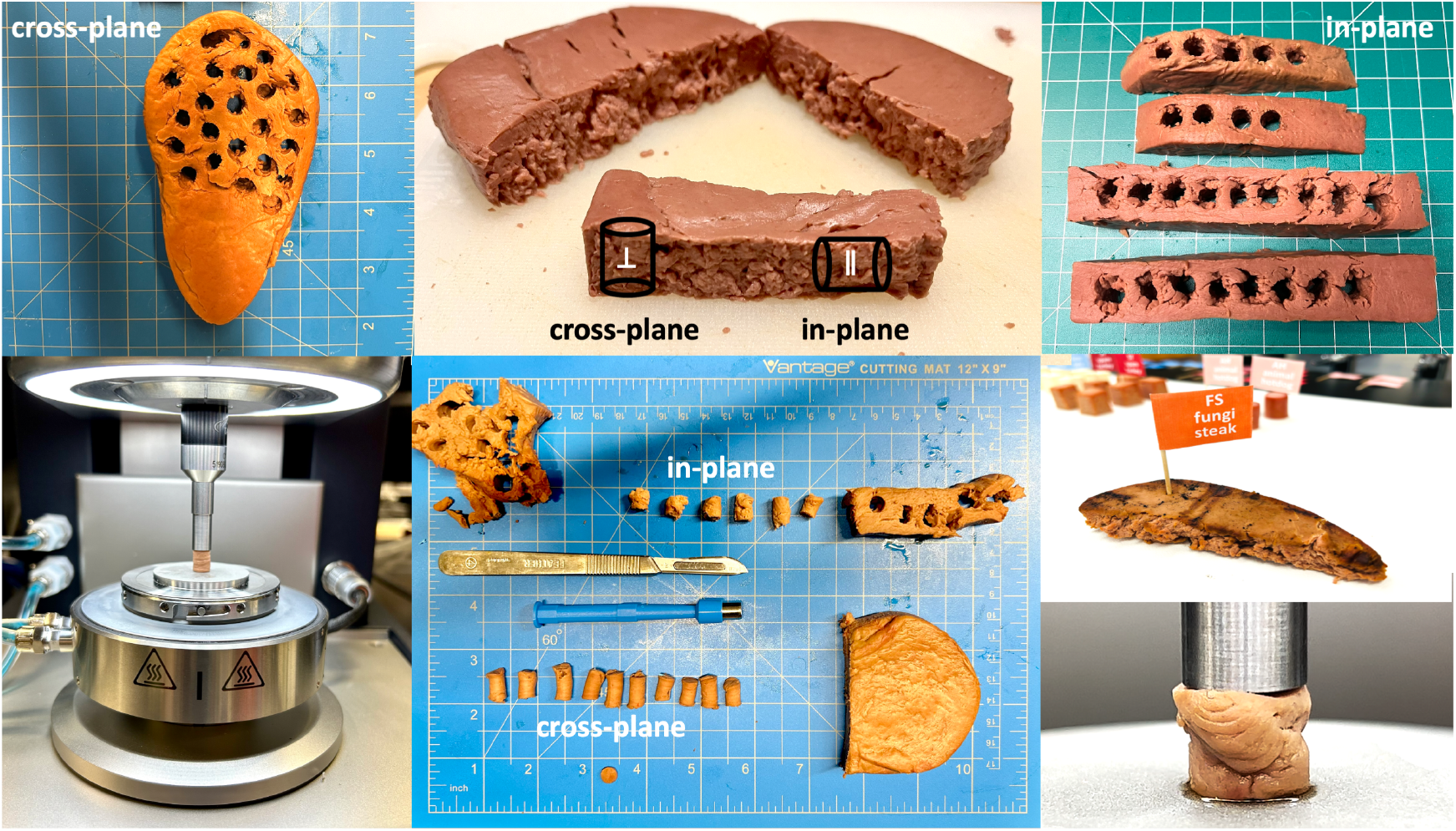
Sample preparation. We test fungi-based meat in compression and shear. To prepare the samples, we use a biopsy punch and extract cylindrical samples with 8 mm diameter and 10 mm height, both in-plane and cross-plane. We test a total of *n* = 164 samples, *n* = 82 in each direction, *n* = 20 in quasi-statically in shear and compression, *n* = 50 in uniaxial double compression at five different frequencies, and *n* = 12 in oscillatory shear.

### 2.2. Sample testing

For each direction, in-plane and cross-plane, we prepare *n* = 20 samples for quasi-static shear and compression testing with ten samples each, *n* = 50 samples for uniaxial double compression testing at five frequencies with ten samples each, and *n* = 12 samples for oscillatory shear testing with two samples for amplitude sweeps and ten for frequency sweeps. We perform all tests using an HR20 discovery hybrid rheometer (TA Instruments, New Castle, DE) and mount the samples between a 40 mm diameter base plate and a 8 mm diameter parallel plate, both sandblasted to avoid slippage. Figure 1 displays our compression and shear test setup with the sample mounted in the rheometer. We guarantee a consistent mounting force below 0.05 N across all samples.

#### 2.2.1. Compression and shear testing for automated model discovery and mechanical analysis

For the quasi-static compression tests, we mount the samples and compress them to a stretch of λ = 0.8 at a stretch rate of 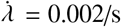, which translates to a total loading time of *t* = 100 s. For the quasi-static shear tests, we mount the samples, fix them at λ = 0.9 compression, and then shear them to a shear strain of γ = 0.1 at a shear rate of 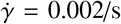, which translates to a total loading time of *t* = 50 s. Figure 2, top left, illustrates the shear stress-strain and compression stress-stretch behavior, where the curves and the shaded regions represent the mean and standard error of the mean across *n* = 10 tests in the in-plane direction parallel to the fibers in orange and in the cross-plane direction perpendicular to the fibers in brown. The quasi-static compression and shear tests serve as the basis for the nonlinear automated model discovery in Section 2.3 and for the linear mechanical analysis in Section 2.4.

**Figure 2:**
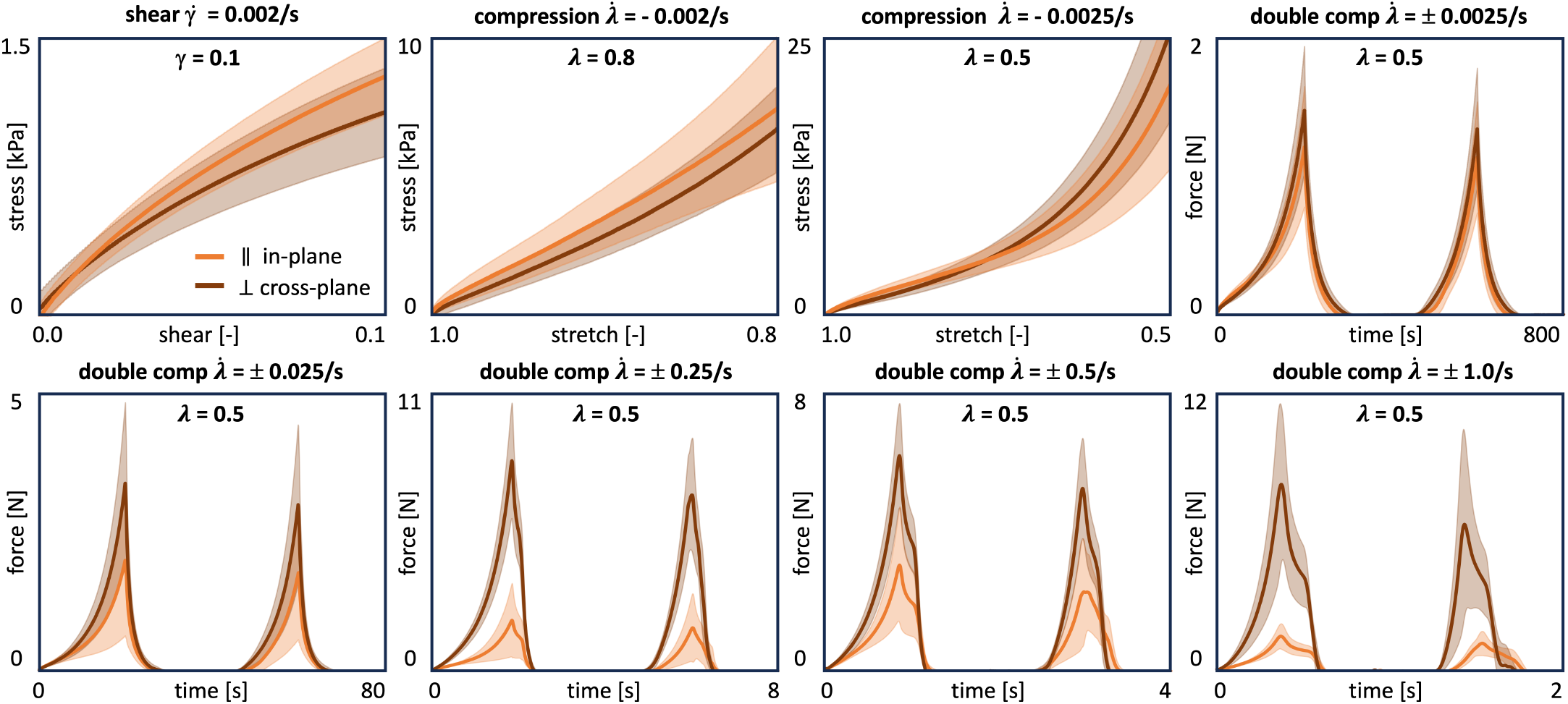
Shear, compression, and double compression testing. Quasi-static shear stress-strain data for γ = 0.1 at 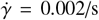 for 50 s, quasi-static compression stress-stretch data for λ = 0.2 at 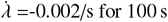 for 100 s, and λ = 0.5 at 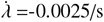 for 200s, and double compression force-time data for λ = 0.5 at five different rates 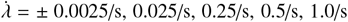 for 800s, 80s, 8s, 4s, 2s. Curves and shaded regions represent the mean and standard error of the mean across *n* = 10 tests in the in-plane direction in orange and the cross-plane direction in brown.

#### 2.2.2. Double compression testing for texture profile analysis

For the double compression tests, we mount the samples, compress them to λ = 0.5 at a rate 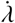, unload the samples at the same rate, and repeat this loading and unloading process a second time. We apply five different rates 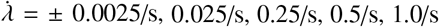 resulting in total test times of 800s, 80s, 8s, 4s, 2s. Figure 2, top right and bottom, illustrates the double compression force-time behavior, where the curves and the shaded regions represent the mean and standard error of the mean of *n* = 10 tests in the in-plane direction parallel to the fibers in orange and in the cross-plane direction perpendicular to the fibers in brown. The double compression tests serve as the basis for the texture profile analysis in Section 2.5.

#### 2.2.3. Oscillatory shear testing for rheological analysis

For the oscillatory shear tests, we mount the samples and fix them at λ = 0.9 compression. Under this pre-load, we first perform *n* = 2 amplitude sweep tests, oscillating at 0.5 Hz, with shear amplitudes increasing logarithmically from γ = 0.0001 to γ = 0.6. Figure 3, top row, shows the recordings of the amplitude sweep, from which we select the shear amplitude of γ = 0.02 to define the linear regime for the subsequent frequency sweeps. We then perform *n* = 10 frequency sweep tests, around γ = 0.02, with angular frequencies increasing logarithmically from ω = 0.1 rad/s to ω = 100 rad/s. Figure 3, bottom row, shows the recordings of the frequency sweep, from which we extract the storage modulus *G*^′^, loss modulus *G*^′′^, complex shear modulus *G*^*^, and phase angle δ, from the data below an angular frequency of ω < 5 rad/s. The oscillatory shear tests serve as the basis for the rheological analysis in Section 2.6.

**Figure 3:**
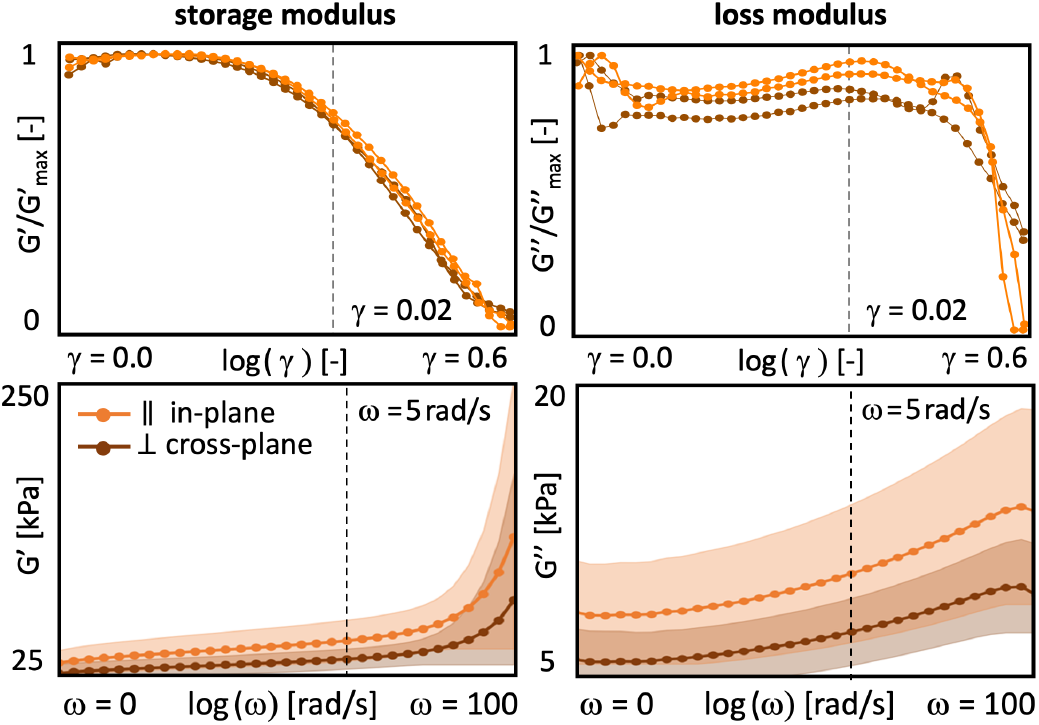
Rheological analysis. The amplitude sweeps, top, oscillate at 0.5 Hz with shear amplitudes increasing logarithmically from γ = 0.0001 to γ = 0.6. We select the amplitude of γ = 0.02 to define the linear regime for the subsequent frequency sweeps. The frequency sweeps, bottom, oscillate around γ = 0.02 with angular frequencies increasing logarithmically from ω = 0.1 rad/s to ω = 100 rad/s. Curves of the amplitude sweep, top, represent the *n* = 2 tests, curves and shaded regions of the frequency sweep, bottom represent the mean and standard error of the mean across *n* = 10 tests with the in-plane direction in orange and cross-plane direction in brown.

### 2.3. Automated model discovery

To characterize the *non-linear elastic* behavior, we perform automated model discovery to discover the *one-term model* and *two-term model* that best explain the quasi-static behavior throughout 50% compression in the in-plane and cross-plane directions. Figure 12 in the Appendix shows our constitutive neural network with its input, its hidden layers, its weights, and its output. Specifically, we use an eight-term network that takes the first and second invariants as input and has two hidden layers with linear and quadratic functions in the first layer and the identity and exponential functions in the second layer [14]. We combine the network with L_1_-regularization using a regularization parameter of α = 0.001 [16], to prevent overfitting, reduce the number of terms to either one or two, and discover the best one- and two-term models out of eight and 28 possible models [17].

### 2.4. Mechanical analysis

To characterize the *linear elastic* behavior, we perform a linear regression on the quasi-static compression and shear data throughout 20% compression and 10% shear to extract the compressive, shear, and mean stiffnesses [11]. We postulate a linear stress-strain relation, σ = *E* · ε, and determine the *compressive stiffness, E*_com_ = (ε · σ)/(ε · ε), from the recorded strain-stress pairs ε; σ using linear regression. Similarly, we postulate a linear shear stress-strain relation, τ = µ · γ, convert the shear modulus µ into the shear stiffness, *E*_shr_ = 2 [1 + ν] µ = 3 µ, and determine the *shear stiffness*, 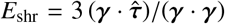, from the recorded shear strain-stress pairs {γ; τ}. Finally, we average the stiffnesses in compression and shear to obtain the *mean stiffness, E*_mean_ = (*E*_com_ + *E*_shr_)/2.

### 2.5. Texture profile analysis

To characterize the behavior during chewing, we perform a texture profile analysis using the double compression tests from Section 2.2.2. Figure 2, top right and bottom, illustrates the load profile of the double compression tests, which consist of two consecutive cycles of 50% compression. Figure 4 shows snapshots of the unloaded and maximally loaded samples for four different loading rates 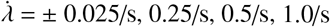. According to standard definitions [18, 19], we denote the peak forces of the first and second loading cycles as *F*_1_ and *F*_2_, the associated loading times as *t*_1_ and *t*_2_, the areas under their loading paths as *A*_1_ and *A*_3_, and the areas under their unloading paths as *A*_2_ and *A*_4_ [13]. We convert the recorded data into stress vs. strain curves, where the stress σ = *F*/*A* is the recorded force *F* divided by the specimen cross section area *A* = π *r*^2^ = 50.3 mm^2^ with *r* = 4 mm, and the strain 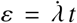 is the applied strain rate multiplied by the time *t* [20]. From the peak force of the first loading cycle *F*_1_, we calculate the peak stress σ_1_ = *F*_1_/*A* at peak strain ε = −50%. From these characteristic values, we extract six texture profile analysis pa-rameters [21]: The *stiffness E* = σ/ε refers to the slope of the stress-strain curve during first compression, when the specimen is compressed to half of its height. The *hardness F*_1_ is associated with the peak force during this first compression cycle. The *cohesiveness* (*A*_3_ + *A*_4_)/(*A*_1_ + *A*_2_) characterizes the material integrity during the second loading and unloading cycle, compared to the first cycle, where a value of one relates to a perfectly intact material, whereas a value of zero relates to complete disintegration. The *springiness t*_2_/*t*_1_ is associated with recovery and viscosity and describes the speed by which the material springs back to its original state after the second cycle compared to the first. The *resilience A*_2_/*A*_1_ is a measure of how well a sample recovers during first unloading path relative to the first loading, where a value of one relates to perfect elasticity whereas a value larger than one indicates plasticity. The *chewiness F*_1_ (*A*_3_ + *A*_4_)/(*A*_1_ + *A*_2_) *t*_2_/*t*_1_, the product of hardness, cohesiveness, and springiness, relates to the resistance of a material during the chewing process, with higher chewiness values indicating that the material is more difficult to chew.

**Figure 4:**
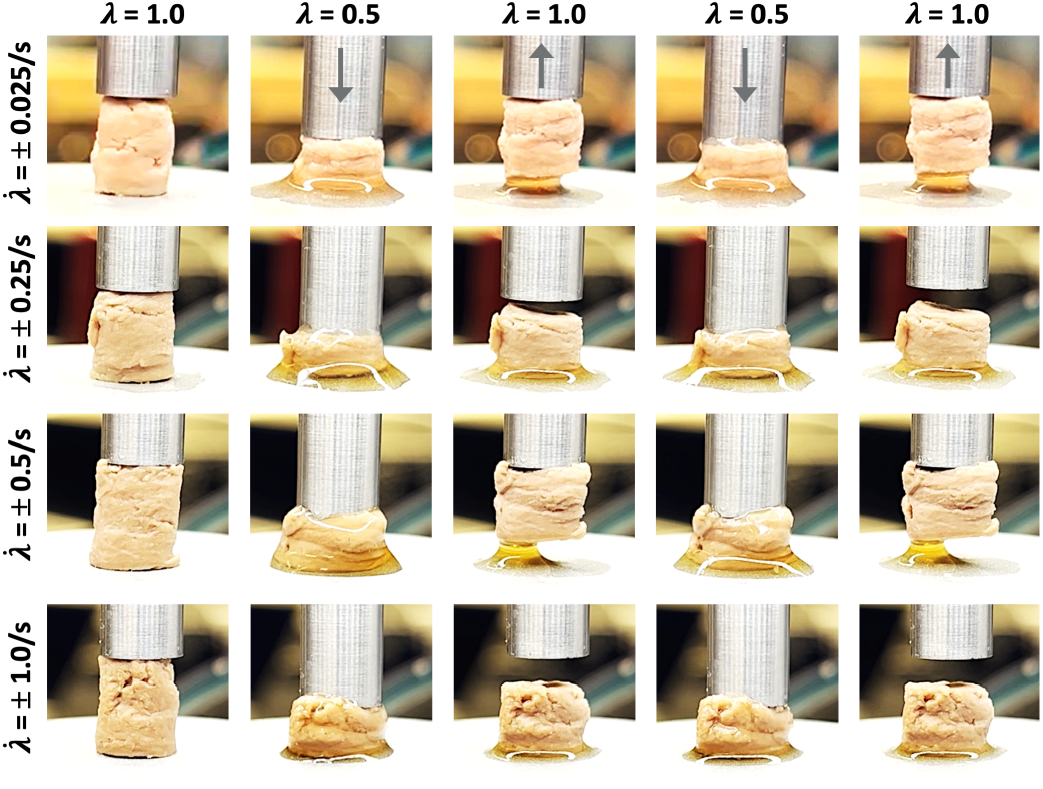
Texture profile analysis. Snapshots of the double compression test showing the unloaded and maximally loaded samples at λ = 1.0, 0.5, 1.0, 0.5, 1.0 for four different loading rates, 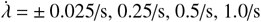, and a total loading time of 80s, 8s, 4s, 2s. The faster the loading, from top to bottom, the less fluid is squeezed out of the sample during the loading process.

### 2.6. Rheological analysis

To characterize the time-dependent behavior, we perform a rheological analysis using the oscillatory shear tests from Section 2.2.3. From the amplitude and frequency sweeps, we extract four rheological parameters [13]: The *storage modulus G*^′^ = *G*^*^ cos(δ) is the apparent or effective stiffness that is a result of both, the stiffness of solid mycelium matrix and the effect of fluid pressure in the mycelial pores. The *loss modulus G*^′′^ = *G*^*^ sin(δ) is the hydraulic dissipation modulus that describes the interstitial fluid flow within the porous mycelium matrix. The *complex shear modulus G*^*^ = *G*^′^+*i G*^′′^ is the sum of the storage and loss moduli and describes the effective poroelastic modulus that combines the elastic and dissipative responses of solid and fluid. The *phase angle* δ = tan^−1^(*G*^′′^/*G*^′^) defines the time lag between stress and strain due to delayed fluid distribution and varies between 0^°^ ≤ δ ≤ 90^°^. The extreme cases of 0^°^ and 90^°^ represent a purely elastic solid that deforms entirely reversibly with all energy stored in the solid mycelium matrix and a purely dissipative fluid that deforms entirely irreversibly with all energy lost to fluid flow.

### 2.7. Sensory survey

For the supplementary sensory survey, we heat the plain fungi-based steaks, without spices or condiments, and prepare bite-sized samples. In accordance with our previous studies, we recruit *n* = 16 participants to participate in the Food Texture Survey [13, 11, 22]. We instruct all participants to eat samples of the product product and rank its texture features on a 5-point Likert scale with twelve questions. Each question starts with “this food is …”, followed by one of the following features [23, 24]: *soft, hard, brittle, chewy, gummy, viscous, springy, sticky, fibrous, fatty, moist*, and *meaty*. The scale ranges from 5 for strongly agree to 1 for strongly disagree. The participants first eat and rank the steak samples in the cross-plane direction, with the fiber direction perpendicular to the chewing direction, and then in the in-plane direction, with the fiber direction parallel to the chewing direction. Between the samples, we ask participants to cleanse their palate with water to minimize residual flavor, neutralize their taste, and standardize the sensory environment. We follow our established protocol and do not include randomization, cross-over design, or sensory blinding to enable consistent comparisons with our previous studies [13, 11, 22]. This research was reviewed and approved by the Institutional Review Board at Stanford University under the protocol IRB-75418.

### 2.8. Statistical analysis

We acquire all data using the TRIOS software and analyze the data sets in Python 3.9. To quantify to which extent the in-plane and cross-plane features differ from each other, we perform a Welch’s t-test for all texture profile analysis, rheological, and sensory features. In the specific texture profile analysis cases of cohesiveness, springiness and resilience, we use a Mann-Whitney U test based on non-continuous data. We report correlations between texture profile analysis features and loading rate using the Spearman and Pearson correlations coefficients ρ and *r*, and correlations between rankings of texture profile analysis and sensory survey features using Kendall’s rank correlation coefficient τ.

## 3. Results

Table 1 provides the detailed information of the product, the Mushroom Root Classic Steak, its brand, ingredients, and nutrition facts. The table also summarizes the discovered one- and two-term models, with their parameters and goodness of fit, and the stiffness features, texture profile analysis features, rheological features, and sensory features. All features are reported in the in-plane and cross-plane directions, with their means and standard deviations. In the following, we compare all features of the fungi-based steak in-plane (FS^||^) and cross-plane (FS^⊥^) against three animal-based products, animal turkey (AT), animal sausage (AS), and animal hotdog (AH), and against five plant-based products, plant turkey (PT), plant sausage (PS), plant hotdog (PH), extrafirm tofu (ET), and firm tofu (FT) [11].

**Table 1:**
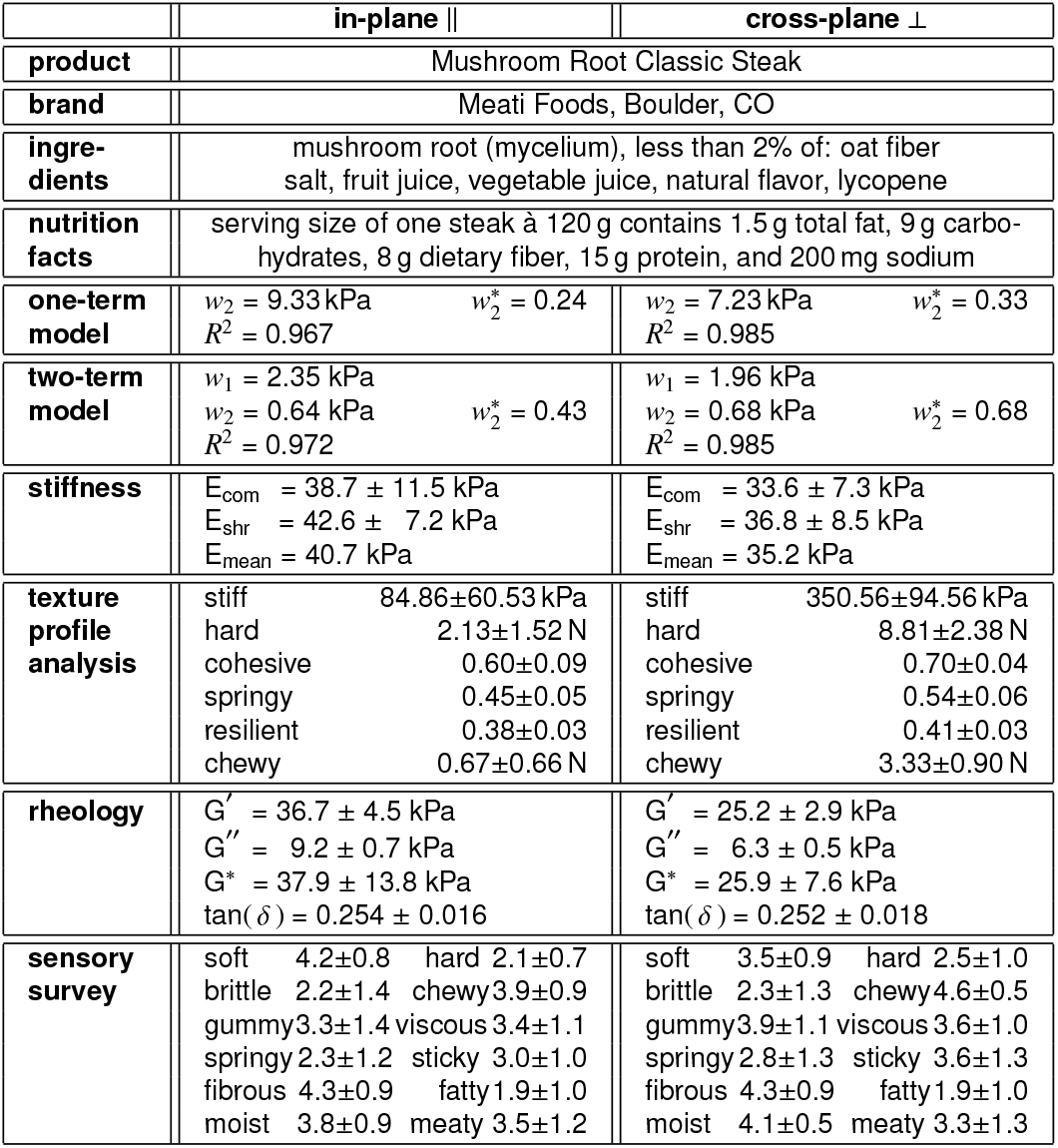
Mushroom Root Classic Steak. Product; brand; ingredients; nutrition facts; discovered one- and two-term models, parameters, and goodness of fit; mechanical features; texture profile analysis features; rheological features; sensory features. All features are reported in in-plane and cross-plane directions with means and standard deviations.

### 3.1. Automated model discovery

Figure 5 summarizes the results of the automated model discovery to characterize the *non-linear elastic* behavior of the fungi-based meat in the in-plane and cross-plane directions for quasi-static compression up to 50%. The dots indicate the experimental data of the *n* = 10 quasi-static compression tests, the dark and light red areas highlight the contributions of the individual model terms, and the *R*^2^ values, the Akaike Information Criterion AIC, and the Bayesian Information Criterion BIC quantify the goodness of fit. Interestingly, the constitutive neural network discovers the same one-term model for both directions, the exponential *Demiray model* [25], 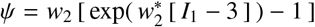, with a linear exponential term of the first invariant, *I*_1_ = λ + 2/λ. Figure 5, top row, shows its fit to the mean experimental data with the discovered stiffness-like and nonlinearity parameters *w*_2_ = 9.33 kPa and 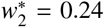 for the in-plane model and *w*_2_ = 7.23 kPa and 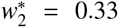 for cross-plane model. Similarly, the network also discovers the same two-term model for both directions, the linear *neo Hookean* model [26] combined with the exponential *Demiray model* [25], 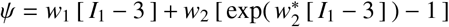, with linear and linear exponential terms of the first invariant. Figure 5, bottom row, shows its fit to the mean experimental data with the discovered stiffness-like and nonlinearity parameters *w*_1_ = 2.35 kPa, *w*_2_ = 0.64 kPa, and 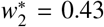 for the in-plane model and *w*_1_ = 1.96 kPa, *w*_2_ = 0.68 kPa, and 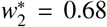 for the cross-plane model. From the *R*^2^ values, the Akaike Information Criterion AIC, and the Bayesian Information Criterion BIC of the one- and two-term models, we conclude that the one-term Demiray model already provides a reasonable first approximation of the data. From the experimental data and from the dominance of the linear neo Hookean term in the small-stretch regime, we conclude that the stretch-stress behavior or the fungi-based meat can be approximated as linear until a compression level of about 80%. This is why we now simplify the analysis and explore the material behavior assuming a linear material response.

**Figure 5:**
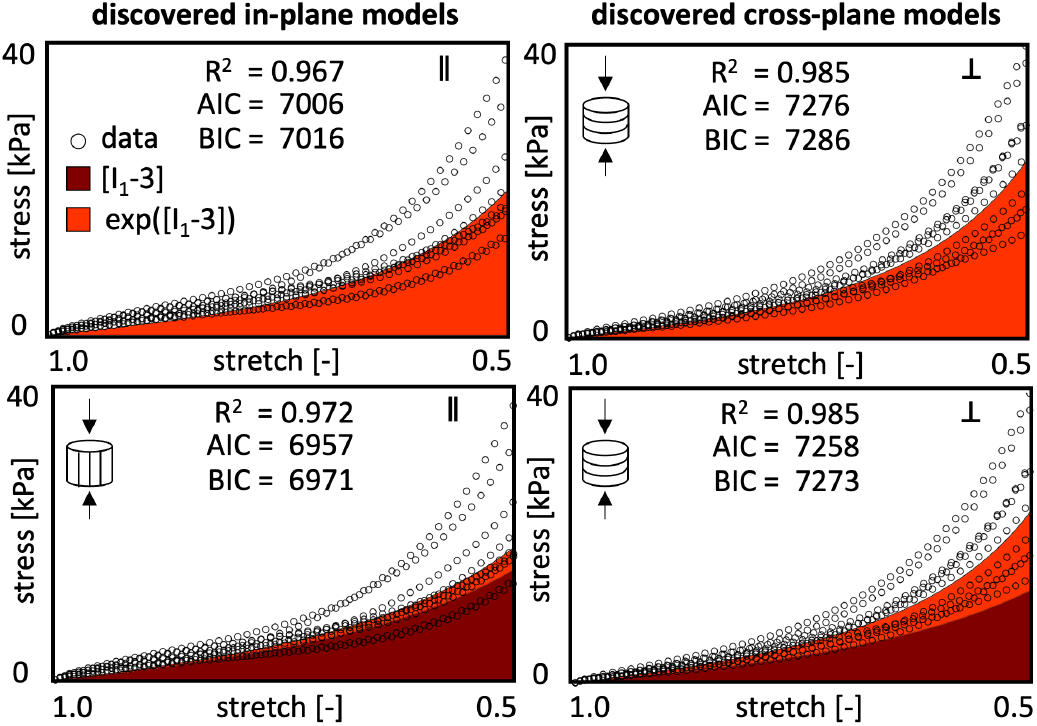
Automated model discovery. Discovered one- and two-term models. The constitutive neural network discovers the exponential Demiray model as the best-fit one-term model and a combination of the linear neo Hookean model and the exponential Demiray model as the best-fit two-term model for both the in-plane and cross-plane directions. The dots indicate the experimental data of the *n* = 10 quasi-static compression tests, the dark and light red areas highlight the contributions of the linear and exponential first-invariant terms; the *R*^2^ values, the Akaike Information Criterion AIC, and the Bayesian Information Criterion BIC quantify the goodness of fit.

### 3.2. Mechanical analysis

Figure 6 summarizes the results of the linear regression to characterize the *linear elastic* behavior of the fungi-based meat in the in-plane and cross-plane directions for quasi-static compression up to 20% and shear up to 10%. The three bar graphs illustrate the in-plane and cross-plane compressive stiffness, shear stiffness, and mean stiffness of the fungi-based steak. All stiffness values are fairly similar, with the stiffnesses in the in-plane direction, E_com_ = 38.7 ± 11.5 kPa, E_shr_ = 42.6 ± 7.2 kPa, and E_mean_ = 40.7 kPa being slightly larger than in the cross-plane direction, E_com_ = 33.6 ± 7.3 kPa, E_shr_ = 36.8 ± 8.5 kPa, and E_mean_ = 35.2 kPa. For comparison, we also include the values of three animal-based products, animal turkey, animal sausage, and animal hotdog, and five plant-based products, plant turkey, plant sausage, plant hotdog, extrafirm tofu and firm tofu [11]. We observe that plant hotdog with 38.2 kPa, animal sausage with 37.5 kPa, and animal hotdog with 37.1 kPa have similar mean stiffnesses as fungi steak, while plant turkey with 223.7 kPa is about five times as stiff, plant sausage with 103.9 kPa and animal turkey with 71.1 kPa are about twice as stiff, and firm tofu with 26.3 kPa and extrafirm tofu with 24.3 kPa are slightly softer.

**Figure 6:**
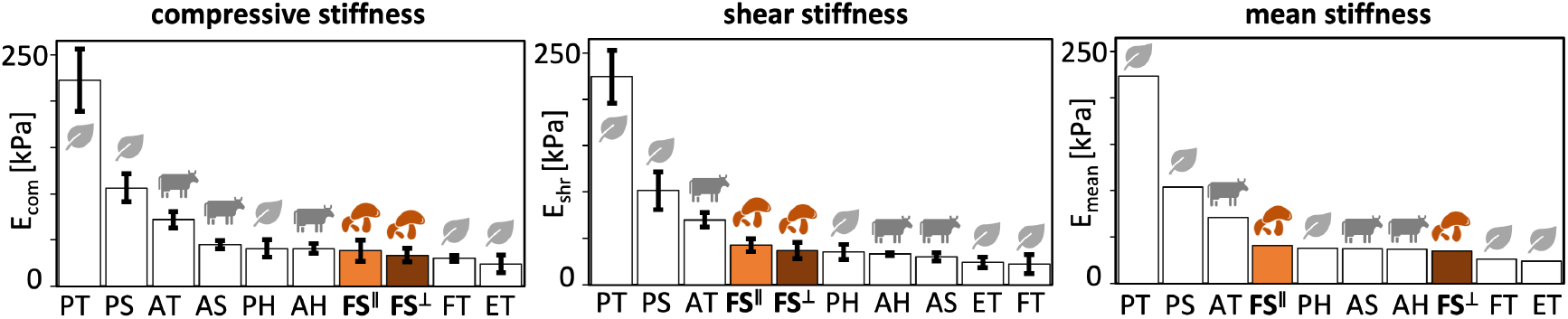
Mechanical analysis. Compressive stiffness, shear stiffness, and mean stiffness. In-plane and cross-plane values for fungi-based product, highlighted in orange and brown, compared to animal- and plant-based products. FS^||^ fungi steak in-plane, FS^⊥^ fungi steak cross-plane, AT animal turkey, AS animal sausage, AH animal hotdog, PT plant turkey, PS plant sausage, PH plant hotdog, ET extrafirm tofu, and FTfirm tofu.

### 3.3. Texture profile analysis

Figure 7 summarizes the results of the double compression test of the fungi-based meat in the in-plane and cross-plane directions for varying loading rates for compression up to 50%. The darkest color highlights the slowest loading rate of 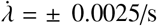 for a total experimental time of 800 s and the lightest color highlights the fastest rate of 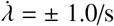 for a time of 2 s. The curves reflect two general trends: The faster the loading, from dark to light, the larger the peak force at 50% compression. At faster loading rates, the cross-plane peak force is about four times larger than the in-plane peak force, while at slow loading rates, the peak forces are almost identical.

**Figure 7:**
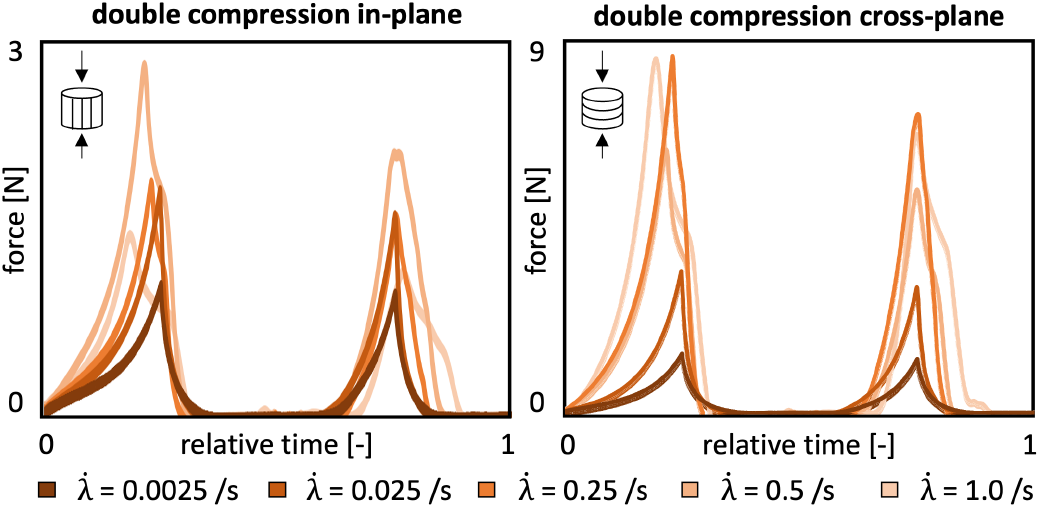
Texture profile analysis. Loading rate sensitivity of force-time response. In-plane and cross-plane force-time behavior during the double compression test for five different loading rates, 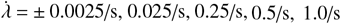, and a total loading time of 800s, 80s, 8s, 4s, 2s. The faster the loading, from dark to light, the larger the peak force at 50% compression.

Figure 8 illustrates the six characteristic features of the double compression test of the fungi-based meat in the in-plane and cross-plane directions for varying loading rates for compression up to 50%. The *resilience, A*_2_/*A*_1_, a measure of how well the sample recovers during first unloading path relative to the first loading path, displays the strongest correlation to the loading rate 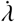. Its Spearman correlation coefficients of ρ = 0.90 (*p* = 0.037) in-plane and ρ = 0.90 (*p* = 0.037) cross-plane indicate a strong, statistically significant, monotonic correlation, and its Pearson correlation coefficients of *r* = 0.996 (*p* = 0.00031) in-plane and *r* = 0.996 (*p* = 0.00052) cross-plane suggest an extremely strong, statistically significant, linear correlation. The *stiffness, E* = σ/ε, the slope of the stress-strain curve during the first compression, and the *hardness, F*_1_, the peak force during this first compression cycle, only display correlations to the loading rate 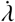 cross-plane, but not in-plane. Cross-plane, its Spearman correlation coefficient of ρ = 0.90 (*p* = 0.037) indicates a strong, statistically significant, monotonic relationship and its Pearson correlation coefficient of *r* = 0.76 (*p* = 0.13) suggests a linear, but not statistically significant trend. Similarly, the *springiness, t*_2_/*t*_1_, the relative time by which the sample recovers, only displays correlations to the loading rate 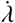 cross-plane, but not in-plane. Cross-plane, its Spearman correlation coefficient of ρ = −0.90 (*p* = 0.037) indicates a strong, statistically significant, negative monotonic correlation and its Pearson correlation coefficient of *r* = −0.944 (*p* = 0.0159) suggests a strong, statistically significant, negative linear correlation cross-plane. The *cohesiveness*, (*A*_3_ + *A*_4_)/(*A*_1_ + *A*_2_), the material integrity during the second cycle, compared to the first cycle, and the *chewiness, F*_1_ (*A*_3_ + *A*_4_)/(*A*_1_ + *A*_2_) *t*_2_/*t*_1_, the product of hardness, cohesiveness, and springiness, do not display a statistically significant correlation or general trend with changing loading rates 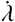 In the following, we focus on the moderate loading rate of 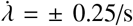, with a total time of 8 s for two complete chewing cycles.

**Figure 8:**
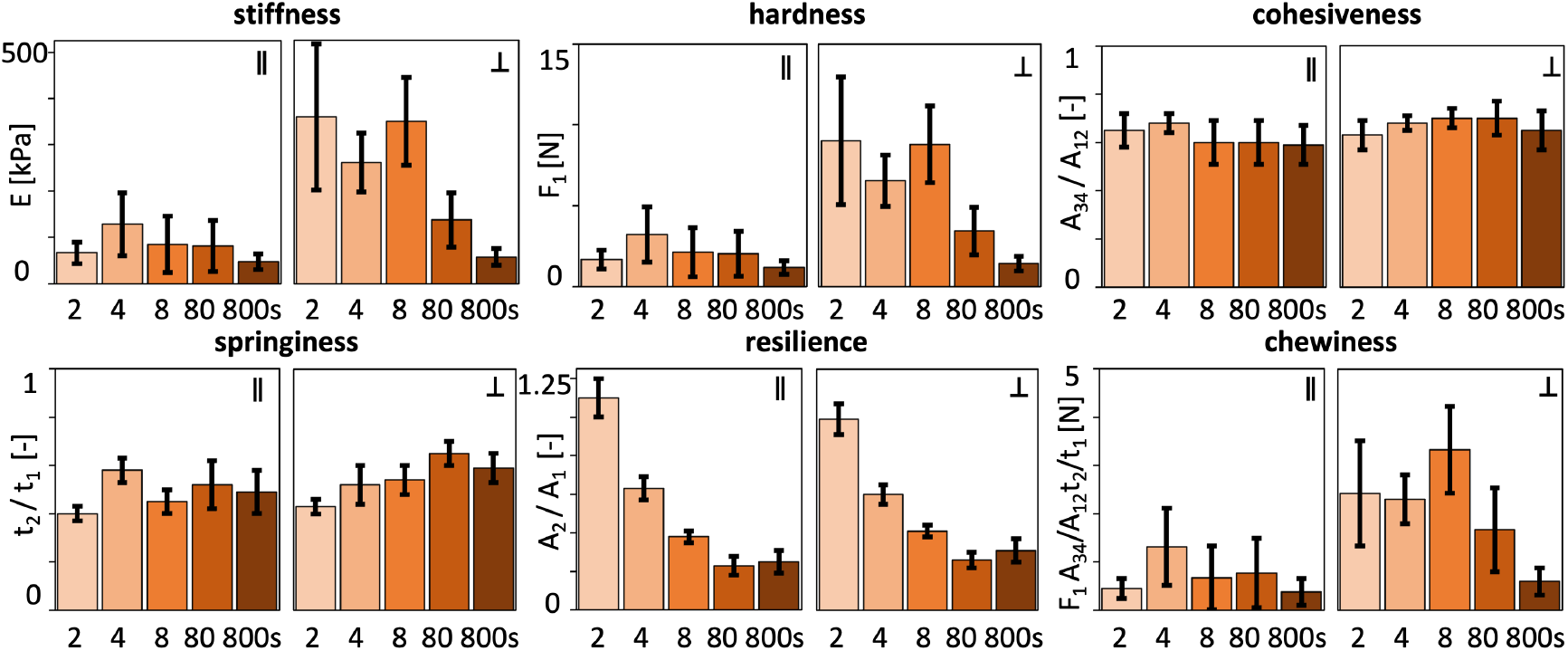
Texture profile analysis. Loading rate sensitivity of six characteristic features. In-plane and cross-plane stiffness, hardness, cohesiveness, springiness, resilience, and chewiness for five different loading rates, 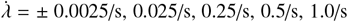, and a total loading time of 800s, 80s, 8s, 4s, 2s. The faster the loading, from dark to light, the larger the stiffness, hardness, and the resilience, and the smaller the springiness.

Figure 9 summarizes the results of the texture profile analysis of the fungi-based meat in the in-plane and cross-plane directions for a double compression to 50% at a loading rate of 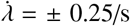. The six bar graphs illustrate the in-plane and cross-plane stiffness, hardness, cohesiveness, springiness, resilience, and chewiness of the fungi-based steak, corresponding to the bright orange bars in Figure 8. Interestingly, for all six features, the cross-plane values are markedly larger than the in-plane values, with statistically significant differences for all features except for resilience. Most notable, the off-plane and in-plane *stiffness E* = 350.56 ± 94.56 kPa and *E* = 84.86 ± 60.53 kPa and the the off-plane and in-plane *hardness F*_1_ = 8.81 ± 2.38 N and *F*_1_ = 2.13 ± 1.52 N vary by more than a factor four, a difference that naturally carries over to the *chewiness* of 3.33 ± 0.90 N and 0.67 ± 0.66 N. At the same time, the *cohesiveness* of 0.70 ± 0.04 and 0.60 ± 0.09, the *springiness* of 0.54 ± 0.06 and 0.45 ± 0.05, the *resilience* of 0.41 0.03 and 0.38 0.03 display only small variations in the off-plane and in-plane directions. Strikingly, the stiffness, hardness, cohesiveness, and chewiness of our fungi-based product all lie within the range of our animal- and plant-based products [13], while its springiness and resilience are significantly smaller both in-plane and cross-plane.

**Figure 9:**
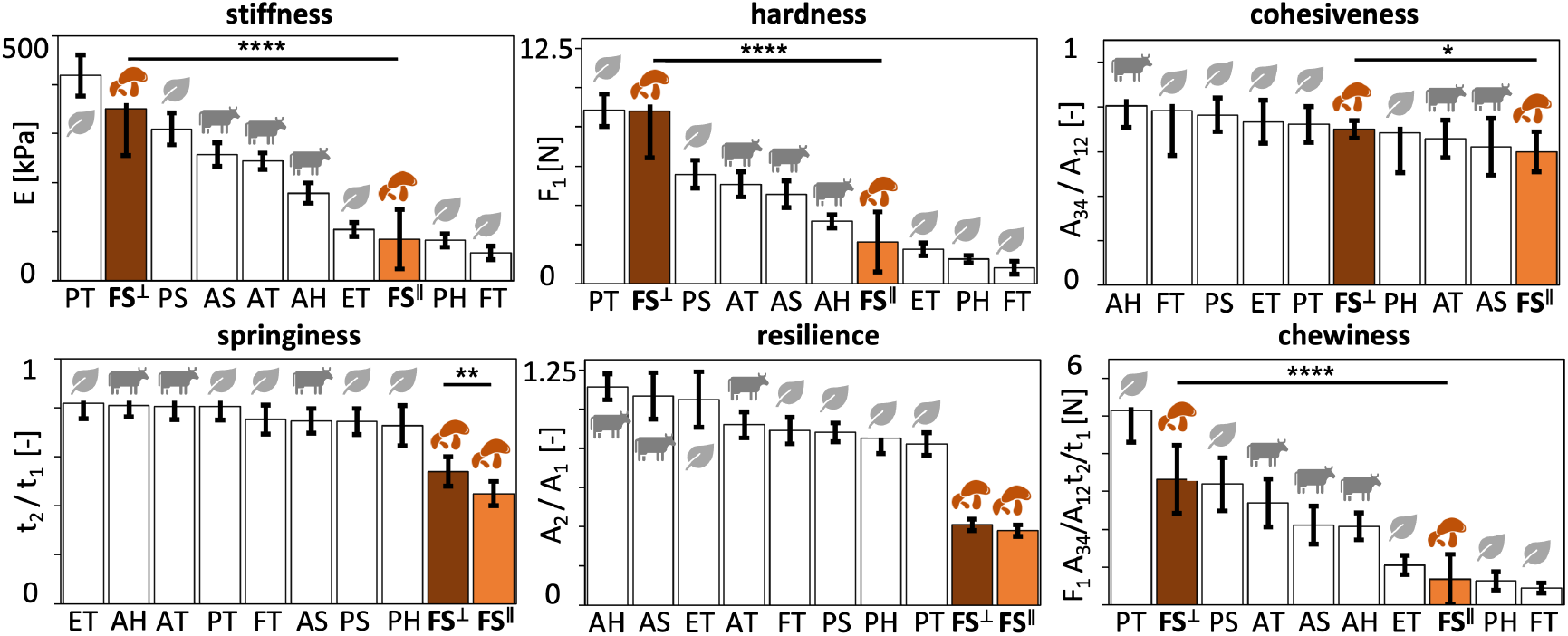
Texture profile analysis. Stiffness, hardness, cohesiveness, springiness, resilience, and chewiness. In-plane and cross-plane values at a loading rate 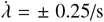, for a total time of 8 s, for fungi-based product, highlighted in orange and brown, compared to animal- and plant-based products. FS^||^ fungi steak in-plane, FS^⊥^ fungi steak cross-plane, AT animal turkey, AS animal sausage, AH animal hotdog, PT plant turkey, PS plant sausage, PH plant hotdog, ET extrafirm tofu, and FT firm tofu. Statistical significance is indicated as * for *p* ≤ 0.05, ** for *p* ≤ 0.01, and **** for *p* ≤ 0.0001.

### 3.4. Rheological analysis

Figure 10 summarizes the results of the rheological analysis of the fungi-based meat in the in-plane and cross-plane directions. The four bar graphs illustrate the in-plane and cross-plane storage moduli, loss moduli, complex shear moduli, and phase angles, of our fungi-based product compared to the animal- and plant-based products [13]. We recorded in-plane and cross-plane storage moduli of *G*^′^ = 36.7 ± 4.5 kPa and *G*^′^ = 25.2 ± 2.9 kPa, loss moduli of *G*^′′^ = 9.2 ± 0.7 kPa and *G*^′′^ = 6.3 ± 0.5 kPa, complex shear moduli of *G*^*^ = 37.9 ± 13.8 kPa and *G*^*^ = 25.9 ± 7.6 kPa, and phase angles of tan(δ) = 0.254 ± 0.016 and tan(δ) = 0.252 ± 0.018. The storage moduli of *G*^′^ = 36.7 kPa and 25.2 kPa quantify the elastic or recoverable part of the mechanical response and suggest that fungi-based steak is stiffer and more elastic in-plane than cross-plane. The loss moduli of *G*^′′^ = 9.2 kPa and 6.3 kPa, quantify the hydraulic or dissipative part of the mechanical response and suggest that fungi-based steak dissipates more energy in-plane than cross-plane, although both values are relatively low compared to the storage moduli *G*^′^. The phase angles of tan(δ) = 0.254 and 0.252 measure the lag between stress and strain and suggest a weak damping and high recoverability both in-plane and cross-plane. These results agree with the our mechanical analysis in the *linear* regime in Figure 6, where the stiffness values in the in-plane direction are slightly larger than in the cross-plane direction. Interestingly, the storage, loss, and complex shear moduli of our fungi-based product all lie well within the range of our animal- and plant-based products. The phase angles are notably larger suggesting a more dissipative, time-sensitive response, likely triggered by the high water content.

**Figure 10:**
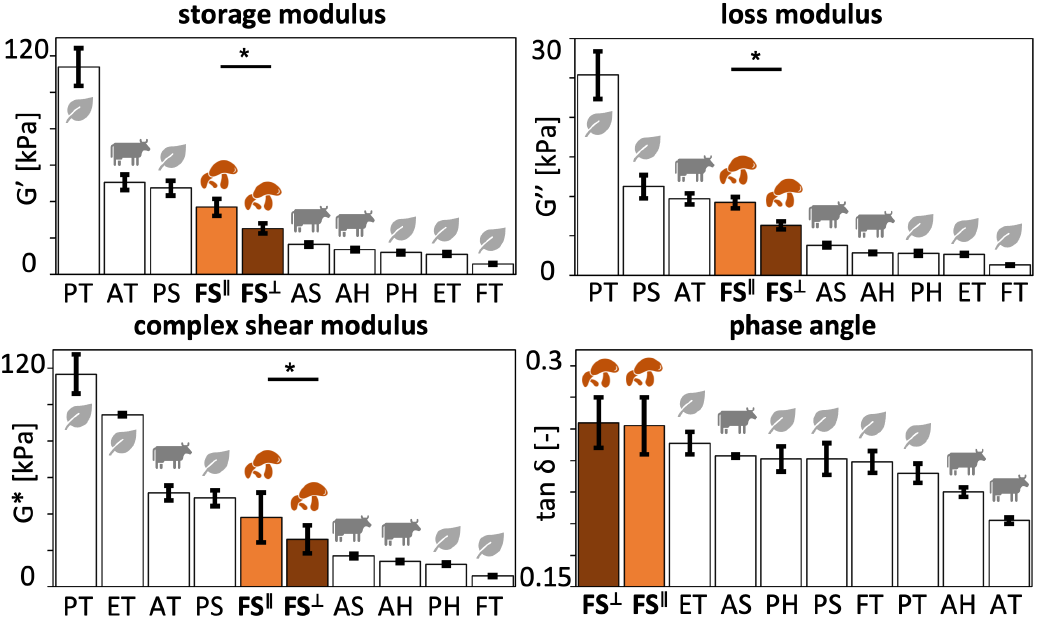
Rheological analysis. Storage modulus, loss modulus, complex shear modulus, and phase angle. In-plane and cross-plane values for fungi-based product, highlighted in orange and brown, compared to animal- and plant-based products. FS^||^ fungi steak in-plane, FS^⊥^ fungi steak cross-plane, AT animal turkey, AS animal sausage, AH animal hotdog, PT plant turkey, PS plant sausage, PH plant hotdog, ET extrafirm tofu, and FT firm tofu. Statistical significance is indicated as * for *p* ≤ 0.05.

### 3.5. Sensory survey

Figure 11 summarizes the results of the sensory survey of the fungi-based meat in the in-plane and cross-plane directions. The twelve bar graphs illustrate the in-plane and cross-plane softness, hardness, brittleness, chewiness, gumminess, viscosity, springiness, stickiness, fibrousness, fattiness, moistness, and meatiness of our fungi-based product compared to the animal- and plant-based products [11]. In the comparison of both directions, the cross-plane scores are larger than the in-plane scores in all categories except for softness and meatiness. In the comparison with the animal- and plant-based products, the fungi-based product stands out in several extremes: It scores highest in five of the twelve categories, *chewiness, viscosity, stickiness, fibrousness*, and *moistness* both in-plane and cross-plane. Both directions also score within the top three in *gumminess* and *meatiness*, and within the bottom three in *springiness*. In the remaining four categories, *softness, hardness, brittleness*, and *fattiness*, the two fungi-based directions score well within the animal- and plant-based products.

**Figure 11:**
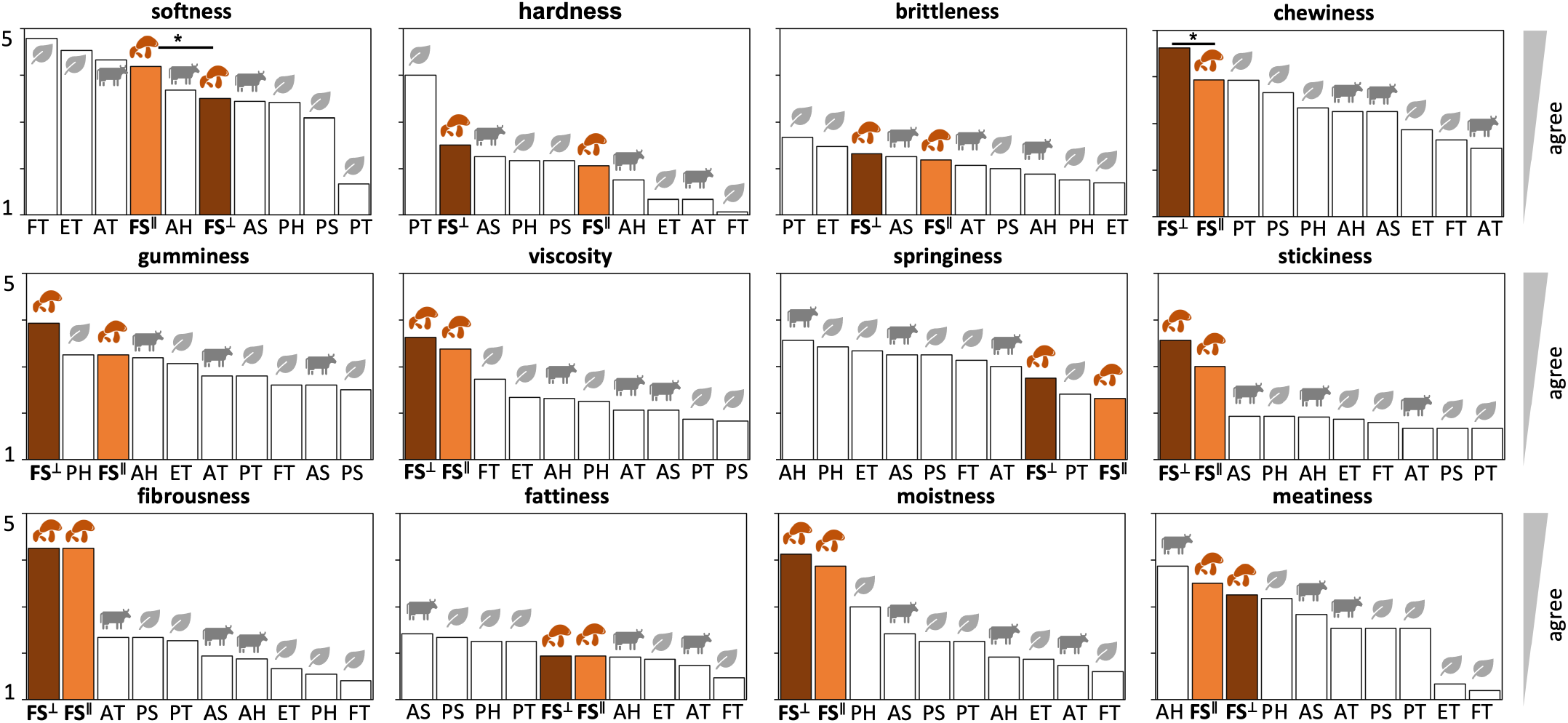
Sensory survey. Softness, hardness, brittleness, chewiness, gumminess, viscosity, springiness, stickiness, fibrousness, fattiness, moistness, and meatiness. In-plane and cross-plane values for fungi-based product, highlighted in orange and brown, compared to animal- and plant-based products. FS^||^ fungi steak in-plane, FS^⊥^ fungi steak cross-plane, AT animal turkey, AS animal sausage, AH animal hotdog, PT plant turkey, PS plant sausage, PH plant hotdog, ET extrafirm tofu, and FTfirm tofu. Statistical significance is indicated as * for *p* ≤ 0.05.

**Figure 12:**
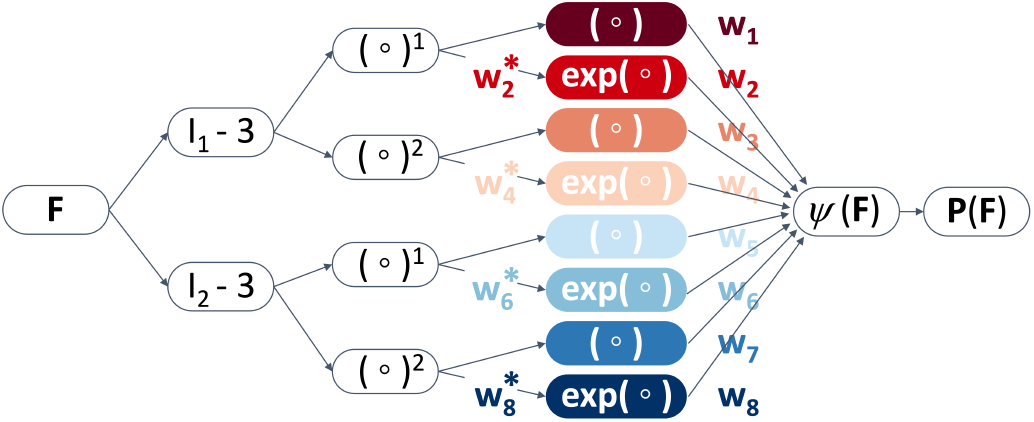
Automated model discovery. The constitutive neural network takes the deformation gradient ***F*** as input and outputs the free energy function *ψ* from which we calculate the stress ***P*** = *∂ψ/∂****F***. The network first calculates functions of the invariants, *I*_1_ and *I*_2_, and then feeds them into its two hidden layers. The first layer generates the first and second powers, (°)^1^ and (°)^2^, of the invariants and weights the even terms by four weights, 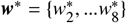. The second layer applies the identity and the exponential function, (°) and exp(°), to these powers. The free energy function *ψ* is the sum of the resulting color-coded terms, multiplied by the eight weights ***w*** = {*w*_1_, …*w*_8_}. We train the network by minimizing the error between model ***P***(***F, w, w***^*^) and data 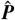 to learn the network parameters ***w*** and ***w***^*^, and apply L_1_ regularization to fine-tune the sparsity of the parameter vector ***w***.

Interestingly, the *softness* rankings from our sensory survey in Figure 11 correlate negatively with the mean *stiffness* rankings from the mechanical analysis in Figure 6, with Kendall’s rank correlation coefficient of τ = −0.51 (*p* = 0.047), while we observed no statistically significant correlation between the sensory *hardness* and mechanical *stiffness*. Strikingly, the *softness* and *hardness* rankings from our sensory survey in Figure 11 correlate negatively and positively with the *stiffness* and *hardness* rankings from the texture profile analysis in Figure Their Kendall’s rank correlation coefficients of τ = − 0.51 (*p* = 0.047) for softness and τ = +0.51 (*p* = 0.047) for hardness indicate a statistically significant correlation. The *chewiness* and *springiness* rankings from the sensory survey and the texture profile analysis are only weakly correlated, with τ = +0.29 (*p* = 0.293) for both features, indicating that the correlation is not statistically significant. Finally, the *viscosity* rankings from our sensory survey in Figure 11 and the *phase angle* rankings from our rheological analysis in Figure 10 display a similar trend in rankings, with τ = 0.42 (*p* = 0.108), but their correlation is not statistically significant.

## 4. Discussion

### Texture profile analysis is sensitive to the loading rate

Figure 7 suggests that, the faster the loading, the larger the peak force. Over a range of two orders of magnitude, for loading rates from 0.0025/s to 0.25/s, the in-plane peak force varies by a factor 1.79, from 1.19N to 2.13N, while the cross-plane peak force varies by a factor 6.08, from 1.45N to 8.81N. Figure 8 confirms that faster loading rates result in larger *stiffness, hardness*, and *resilience* values, and in smaller *springiness* values. A closer look at the double compression test in Figure 4 provides an explanation for these observations: At slow loading rates, the fluid has enough time to gradually escape the sample. In the extreme case of an infinitely slow loading rate, the machine only probes the solid mycelium matrix. At fast loading rates, the fluid remains trapped in the sample. In the extreme case of an infinitely fast loading rate, the machine probes both the solid mycelium matrix and the enclosed fluid. The classical texture profile analysis is primarily a mechanical test that simulates the action of chewing; it focuses on elastic and plastic deformations, but not on dissipative behavior in the rheological sense [18]. Yet, our observations confirm that, especially for foods with a high water content, the texture profile analysis is highly sensitive to the magnitude of compression λ and to the loading rate 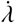 [27]. To allow for comparisons across different studies and laboratories, these parameters should be standardized or, at a very minimum, be carefully reported and discussed along with the results [28]. For a comparison with previous studies, we recommend a compression level of 50% and a loading rate of 0.25/s [13]. Taken together, we need to be extremely careful when comparing the results of texture profile analyses across different studies with varying magnitudes and rates.

### The elastic behavior of fungi-based steak is isotropic under shear and compression

Figure 2 suggests that the quasi-static response of the fungi-based steak to small shear and compression is almost identical in-plane and cross-plane. Interestingly, the shear response softens with increasing deformation, already at 10% shear, first graph, while the compressive response remains relatively linear, even at 20% compression, second graph. Beyond 25% compression, third graph, we observe a significant stiffening with an interesting cross-over, where the cross-plane behavior shown in brown is initially slightly softer, but then stiffens slightly faster than the in-plane behavior shown in orange. Figure 5 confirms these observations with the same discovered one- and two-term models in both directions. The best one-term model is the Demiray model [25] with a linear exponential first invariant term. The discovered parameters feature a slightly lower initial stiffness cross-plane with *w*_2_ = 7.23 kPa compared to in-plane with *w*_2_ = 9.33 kPa, but a faster stiffening cross-plane with 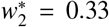 than in-plane with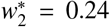. The best two-term model is a combination of the neo Hookean model [26] and the Demiray model [25] with linear and linear exponential first invariant terms. The discovered parameters suggest a dominance of the linear term in-plane with *w*_1_ = 2.35 kPa, illustrated through the large dark red area in Figure 5 bottom right, and an equal contribution of both terms cross-plane with *w*_1_ = 1.96 kPa, illustrated through the smaller dark red area in Figure 5 bottom left. Figure 6 confirms that the quasi-static response to small deformations in shear and compression is almost identical in-plane and cross-plane with compressive, shear, and mean stiffnesses varying moderately between 33.6 kPa and 40.7 kPa, all well within the range of the animal- and plant-based comparison products. At first glance, this observation seems counter-intuitive: Many biomaterials are fibrous and anisotropic in nature [17]. Mushroom root mycelium is composed of a dense, interconnected network of filamentous hyphae, long and tubular structures that branch and fuse to form its complex porous, anisotropic microstructure [7]. Its fibrous matrix is essential for creating its characteristic moisture retention, texture, and mouthfeel that resemble animal meat [2]. It would seem natural to expect that the filamentous hyphae create anisotropic mechanical properties with different porosities and stiffnesses along and across the dominant hyphal orientation. We need to keep in mind though that we have only tested in compression and shear. Most biomaterial models are anisotropic only in tension, and deactivate their fiber contributions during compressive loading [29] which would agree with our present observations. We are currently testing fungi-based steak in tension to explore to which extent its tensile stiffness varies in the in-plane and cross-plane directions. Taken together, we conclude that under quasi-static shear and compression, the in-plane and cross-plane responses of fungi-based steak are not significantly different, suggesting that, to a first approximation, the elastic behavior of fungi-based steak in shear and compression is isotropic.

### The inelastic behavior of fungi-based steak is anisotropic under compression

Figure 2 suggests that the dynamic response of the fungi-based steak to moderate compression is stiffer cross-plane than in-plane. Strikingly, at 50% compression and a loading rate of 0.25/s, the stiffnesses and peak forces of the double compression test vary by a factor four, from 350.56 kPa and 8.81 N cross-plane to 84.86 kPa and 2.13 N in-plane. Figure 9 confirms this observation and shows that, in fact, all cross-plane features of the texture profile analysis are larger than their in-plane counterparts, with statistical significance across all features, stiffness, hardness, cohesiveness, springiness, and chewiness, except for resilience. During the production process, the fungi-based steaks are pressed into shape creating an anisotropic microstructure with a pronounced hyphae orientation in the plane of the steak. We hypothesize that, when loaded in-plane, fluid can escape the sample more easily than when loaded cross-plane. At fast loading rates, this time-dependent effect is dominant, resulting in larger stiffnesses and peak forces cross-plane than in-plane. At slow loading rates, the time-independent elastic is dominant, resulting in more isotropic stiffnesses and peak forces. Our fungi-based steak is made up of 95% mushroom root mycelium with a water content of more than 80% [30]. The large amount of fluid leaving the sample in Figure 4 suggests that a time-dependent poroelastic model [31] would be appropriate to characterize its constitutive behavior across all loading rates [32]. We are currently performing consolidation-type tests [33] to complement our double compression tests at different loading rates and our rheological frequency sweeps to accurately characterize the poroelastic response of a variety of different fungi-based meats. Taken together, we conclude that under dynamic loading, the in-plane and cross-plane responses of fungi-based steak are significantly different, suggesting that the inelastic behavior of fungi-based steak is anisotropic.

### Our sensory perception agrees with the texture profile analysis

Figure 11 suggests that we perceive fungi-based steak as more *chewy*, more *viscous*, more *sticky*, more *fibrous*, and more *moist* than the animal- and plant-based comparison meats. Strikingly, we also perceive it as more *meaty* than all products, except for animal hotdog. In agreement with the texture profile analysis in Figure 9, we perceive the fungi-based steak as less soft, harder, more springy, and more chewy cross-plane than in-plane. In agreement with the rheological analysis in Figure 10, we perceive the fungi-based steak as more dissipative or viscous cross-plane than in-plane. Interestingly, our overall softness and hardness rankings from our sensory survey in Figure 11 display a statistically significant correlation with the stiffness and hardness rankings from the texture profile analysis in Figure 9, demonstrating that we can indeed taste stiffness and hardness [11]. Our observations suggest that we can even perceive small textural differences between the in-plane and cross-plane features of fungi-based steak. Most importantly, we perceive pronounced textural differences between fungi-based steak and the eight animal- and plant-based comparison meats. We need to keep in mind though that not only our plant-based products, but also our animal-based products are processed and isotropic [11]. It would be interesting to compare the fungi steak with an animal steak with a similar fibrous and anisotropic microstructural architecture.

### Fungi-based steak is a viable alternative to animal- and plant-based meats

Compared to animal-based meat, fungi-based steak is healthier for people and for the planet [2]. It has a *high fiber content*, and is rich in beta-glucans and other prebiotic fibers that have a positive impact on our cholesterol levels and gut health [6]. It *grows fast* and sustainably, and can be cultivated within days on vertical farms using agricultural waste or low-cost feedstock [34], with *minimal land and water use* [4]. It has a *low environmental footprint*, and produces fewer greenhouse gases than animal agriculture [35]. Compared to plant-based meats, fungi-based steak has a *lean ingredient list* and is not ultra-processed [36]. Made up of 95% mushroom root mycelium that grows in a filamentous, fibrous structure [7], it has a *natural meat-like texture*, a *natural umami-type flavor*, and a complete amino acid profile [2]. Its is *minimally processed*, it requires no additives, gums, or binders, it is *allergen-friendly*, and free of soy, gluten, and nuts [37]. A recent blinded, randomized tasting study with more than 2,000 participants suggests that fungi- and plant-based steaks generally receive a much lower acceptance rate than animal steaks [38]. Our study demonstrates though that fungi-based steaks can comfortably reproduce the mechanical signature of animal- and plant-based meats: Its *stiffness* and *hardness* lie well within the range of the comparison meats [13], both in-plane and cross-plane. At the same time, its *high viscosity* and *high moisture* outperform all comparison products. Most notably, with only minimal processing, the fungi-based steak achieves a *more fibrous texture* than all eight comparison products. Our texture profile analysis and our sensory survey both confirm that we can reliably and reproducibly measure and sense significant differences in its in-plane and cross-plane behavior. Strikingly, the fungi-based steak also achieves a *more meaty flavor* than seven of the eight comparison products, which supports the general notion of the umami-type taste and meat-like flavor of mushroom root mycelium, its main ingredient [2].

## 5. Conclusion

Fungi-based meats are an emerging class of alternative proteins that offer a nutritious, sustainable, and minimally processed solution to the growing global demand for meat. Our study demonstrates that fungi-based steak–made from 95% of mushroom root mycelium–is a promising, transformative alternative to both animal- and plant-based meats: Under quasistatic shear and compression, its elastic response is isotropic. In contrast, under dynamic compression, its inelastic response is markedly anisotropic—likely as a result of its high moisture content and its inherently fibrous microstructure, with mycelium as its primary ingredient. Our texture profile analysis confirms that the mechanical stiffness, hardness, and resilience of this moist product are highly sensitive to both loading magnitude and rate, highlighting the need for standardized testing protocols. Most notably, the mechanical behavior of fungi-based steak lies well within the range of eight commercial animal- and plant-based meats. Strikingly, in some aspects–especially moisture retention and fibrousness–it outperforms all eight comparison products of our study. Our sensory analysis corroborates these mechanical findings and suggests that fungi-based steak not only mimics a meat-like texture, but also delivers a more umami-rich, meaty flavor than most plant-based alternatives. Taken together, these findings position fungi-based steak as a nutritious, sustainable, and texturally compelling candidate in the future of alternative proteins.

## Data Availability

Data and analysis script are freely available at https://github.com/LivingMatterLab/CANN.

## Acknowledgments

This research was supported by the Research Foundation Flanders FWO through the doctoral fellowship SB1SE2123N to Thibault Vervenne, by the NSF Graduate Research Fellowship, by the Stanford DARE Fellowship, and by seed funding from the Stanford Plant-Based Diet Initiative to Skyler St. Pierre, and by seed funding from Food System Innovations and from the Stanford Doerr School of Sustainability Accelerator, by the NSF CMMI grant 2320933, and by the ERC Advanced Grant 101141626 to Ellen Kuhl. The authors acknowledge the use of ChatGPT, OpenAI, 2025, to assist in generating the graphical abstract.

## Appendix

### 2.3.1. Automated model discovery

We perform automated model discovery to discover the one- and two-term models that best explain the quasi-static behavior in the in-plane and cross-plane directions [17]. Figure 12 shows our eight-term constitutive neural network that takes the first and second invariants, *I*_1_ and *I*_2_, of the deformation gradient ***F*** as input and outputs the free energy function *ψ* from which we calculate the Piola stress, ***P*** = *∂ψ/∂****F*** [39]. Our network has two hidden layers with linear and quadratic functions in the first layer, (°)^1^ and (°)^2^, and the identity and exponential functions in the second layer, (°) and exp(°) [14]. It has four unit-less weights 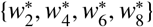 between the first and second layers, and eight weights 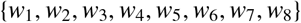 with unit kilopascal out of the second layer. As such, it features a total of twelve network weights that enable 2^8^ combinations of terms resulting in 256 possible models. We learn the networks weights by training the network on the quasi-static compression data up to 50% compression from Figure 2, first row, third graph. We minimize the loss function L that penalizes the error between the model ***P***(***F***_*i*_, ***w, w***^*^) and the data 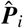, divided by the number of data points *n*_data_. We apply L_1_ regularization or lasso [40], and supplement the loss function by the product of the L_1_ norm of the parameter vector ***w***, weighted by a penalty parameter *α*,

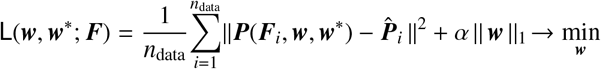

We select a regularization parameter of *α* = 0.001 [16], to prevent overfitting, reduce the number of terms to either one or two, and discover the best one- and two-term models out of eight and 28 possible models [17]. We note that we could have performed the same process by using symbolic regression instead of constitutive neural networks [41].

